# Metabarcoding metacommunities: time, space, and land use interact to structure aquatic macroinvertebrate communities in streams

**DOI:** 10.1101/2022.05.01.490210

**Authors:** Jennifer Erin Gleason, Robert H. Hanner, Karl Cottenie

## Abstract

There is an increasing need to move beyond evaluating the effect of land use on stream communities by only studying local variables, and instead incorporate a metacommunity perspective which integrates environmental and spatial factors across larger spatial scales. The use of molecular tools (DNA metabarcoding) to identify bioindicator groups, such as aquatic macroinvertebrates, can provide greater taxonomic resolution to explore patterns in stream metacommunities. In this study, we collected aquatic macroinvertebrates from streams in southern Ontario which spanned a gradient of agricultural disturbance and used DNA metabarcoding to identify the species composition from these samples. We address a significant knowledge gap in previous stream aquatic macroinvertebrate metacommunity studies by incorporating molecular identification as well as a temporal component. We observed that a combination of local habitat conditions, regional agricultural land use, and spatial position influenced aquatic macroinvertebrate community composition, suggesting there is an interaction between environmental filtering and dispersal processes that structures these communities. However, aquatic macroinvertebrate communities were also highly dissimilar between streams and composed of many rare species, and a large percentage of unexplained variation suggests that there is a strong stochastic component to community assembly. We also observed that there is a seasonal component to metacommunity dynamics, with different water quality variables being significant to community composition in each sampling month. While we expected that an increased percentage of surrounding agricultural land use would result in more homogenous macroinvertebrate communities, we only detected this relationship in May and found evidence that a larger riparian buffer width can mitigate the effects of agricultural land use. We demonstrate the utility of DNA metabarcoding for revealing patterns in metacommunity dynamics that may not be detectable using coarse taxonomic identifications, and reveal the importance of incorporating a seasonal component when evaluating the influence of land use on community composition.

## 1. Introduction

Agricultural land use can alter biological communities in multiple ways, through both local changes in environmental conditions and regional modifications to habitat availability and connectedness. Freshwater ecosystems, such as streams, are sensitive to agricultural impairment, and alterations in habitat quality pose threats to aquatic biodiversity (Albert et al., 2021; Dudgeon, 2019; Reid et al., 2019), as well as the complex multi-trophic food webs that streams support (Champagne et al., 2022; Hladyz et al., 2011). Maintaining the biological integrity of streams is a crucial goal of conservation and restoration programs, as streams provide irreplaceable services, and the maintenance of freshwater ecosystems is vital for both human prosperity and the preservation of biodiversity (Reid et al., 2019). In addition to their extrinsic value, streams not only support local biodiversity (e.g., within the stream channel itself and the adjacent terrestrial habitat), but also transfer resources downstream (i.e., the river continuum concept; Vannote et al., 1980) and into larger water bodies across the landscape (e.g., Wipfli & Gregovich, 2002; Richardson et al., 2021). It is therefore optimal to approach stream conservation using an integrative perspective, which incorporates both local and landscape-scale factors. (Reid et al., 2019). The preservation of freshwater habitats is of paramount importance, and both biomonitoring and ecology have been identified as key research priorities to prevent further loss of these systems (Maasri et al., 2022).

Aquatic macroinvertebrates are important components of stream ecosystems and are often used as bioindicators of environmental condition in freshwater ecosystems due to their functional and taxonomic diversity as well as their documented sensitivities to changes in water quality (Merritt et al., 2008; Rosenberg & Resh, 1993). As streams are closely tied to the terrestrial landscape (see Baxter et al., 2005; Hladyz et al., 2011), the use of aquatic macroinvertebrates is particularly relevant as they link terrestrial and aquatic ecosystems through the emergence of aquatic insect larvae to terrestrial winged adults. Terrestrial changes, such as agricultural land use, can alter stream habitats (and thus influence benthic macroinvertebrates) through changes in water inputs, run-off of pesticides, fertilizers and sediments from adjacent fields, and the alteration of stream morphology (e.g., channelization to increase space for crops; Allan, 2004; Reid et al., 2019). The reduction or removal of riparian zone vegetation reduces habitat complexity, increases erosion, and results in higher water temperatures (Allan, 2004), and can alter stream food webs by removing the input of terrestrial subsidies (i.e., allochthonous nutrient sources; Nakano et al., 1999). In addition to the influence of local and landscape-scale processes described above, it is possible that the effects of agricultural impairment on aquatic systems can vary seasonally because of the variation and timing of insect life cycles in streams (Merritt et al., 2008). The assembly and maintenance of aquatic macroinvertebrate communities are the result of local, regional, and temporal processes, making these systems ideal for the incorporation of metacommunity ecology (Leibold et al., 2004; Leibold & Chase, 2017).

Metacommunity theory is an important concept in modern ecological research and describes several paradigms linking local and regional communities through the dispersal of interacting taxa (Leibold et al., 2004; Leibold & Chase, 2017). Two main themes in metacommunity research are influences of environmental heterogeneity and dispersal processes, building upon the dichotomy of deterministic (i.e., niche) and stochastic (i.e., neutral) processes in ecology (e.g., Chase, 2014; Mikkelson, 2005; Wennekes et al., 2012). Three of the metacommunity structuring processes described by Leibold et al. (2004; species-sorting, patch dynamics, and mass-effects) emphasize the role of environmental filtering (deterministic processes) in structuring communities. The fourth paradigm is the neutral model, where differences among communities are attributed to stochastic or random processes (e.g., immigration, emigration, local extinctions, ecological drift; Chave, 2004; Hubbel, 2001). Adler et al. (2007) suggest that niche and neutral theory should not be considered as competing ideas or mutually exclusive concepts, since both processes can be important in shaping communities. For example, Thompson and Townsend (2006) observed that riverine macroinvertebrate communities were best predicted by a model which combined both local environmental conditions and spatial distance, though this pattern differed for different functional groups. Focusing solely on local factors limits our comprehension of the dynamics and structure of communities (Brown et al., 2011). While environmental factors have been cited as being more important to metacommunity dynamics in smaller, first-order streams (Göthe et al., 2013; Heino et al., 2012), the spatial position of the stream branch is also important in structuring communities (Brown et al., 2017; Grant et al., 2007). The incorporation of metacommunity ecology into bioassessment is a necessary step towards understanding how communities are affected by land use at multiple spatial scales (Heino, 2013).

A limit of previous metacommunity studies on riverine macroinvertebrates is that studies are performed at coarse or mixed taxonomic resolution (e.g., Cañedo-Argüelles et al., 2015; Grönroos et al., 2013; Heino et al., 2004; Ligeiro et al., 2010; Yates & Bailey, 2010) and in some cases exclude difficult-to-identify taxa, such as chironomids (e.g., Astorga et al., 2014). The integration of molecular identification methods is advantageous in both biomonitoring applications and ecological research, particularly for detecting diverse organisms such as invertebrates (Baird & Hajibabaei, 2012; Taberlet et al., 2012). DNA metabarcoding (Taberlet et al., 2012) refers to the sequencing of an entire sample at once and then matching sequences to a reference database (e.g., BOLD – Barcode of Life Data Systems; Ratnasingham & Hebert, 2007) to assign taxonomic information. In a call to action for freshwater preservation, Maasri et al. (2022) identified multiple research priorities, including ecology and biomonitoring, and suggest that these fields can benefit from the incorporation of molecular methods for taxonomic identification. Using metabarcoding, Gleason et al. (2022) discovered that headwater streams in southern Ontario varied substantially in community composition at very small spatial scales (e.g., over ten meters) and that aquatic macroinvertebrate communities were largely comprised of rare taxa. Gleason et al. (2022) detected over 1600 Operational Taxonomic Units (OTUs, a proxy for species) from 149 families, and a single family (Chironomidae) made up one third of the detected OTUs, reflecting the levels of diversity that are not recovered when using family or even genus-level identification. Aquatic macroinvertebrates communities thus contain a high amount of diversity, which is not apparent at coarser taxonomic resolution.

While initial DNA metabarcoding focused primarily on identifying taxonomic composition and richness metrics, this growing field holds great promise for metacommunity ecology (Joly et al., 2014). The potential to process numerous samples efficiently and achieve fine-resolution taxonomic information provides the opportunity for large-scale field studies to shed light on the often-complex interactions of environmental and spatial factors. For example, in a wetland complex in Alberta, Bush et al. (2020) observed highly diverse aquatic macroinvertebrate communities using DNA metabarcoding. The high spatial and temporal turnover could indicate that wetland invertebrate community assembly is nearly random, and Bush et al. (2020) suggested that stochasticity is driving metacommunity dynamics in this system. In other ecosystem types, DNA metabarcoding has been used to explore community assembly and spatial turnover of soil invertebrates (Arribas et al., 2021; Noguerales et al., 2021). In one study, there was very high taxonomic turnover between soil samples, and similarity decreased with spatial distance, suggesting that these communities were predominantly structured by spatial components (Arribas et al., 2021). However, Noguerales et al. (2021) found that both environmental filtering and spatial arrangement were important structuring components of soil invertebrates. These studies illustrate the applicability of DNA metabarcoding for metacommunity ecology, and a key uncertainty which remains is how metacommunity dynamics change in response to agriculture in stream systems where 1) there are no discrete community boundaries and 2) community composition can vary drastically at small spatial scales (Gleason et al., 2022).

In this study, we quantify the relative influence of both regional and local habitat conditions on the community composition of aquatic macroinvertebrates using DNA metabarcoding to gain greater insight into the effects of agriculture on metacommunity dynamics. As stream communities will change seasonally due to differences in hatching and emergence periods, we expect benthic communities will be highly structured by sampling period and that their responses to environmental conditions may not be static over time. We thus test the variation explained by regional agriculture, local habitat parameters, and geographic location during three sampling months (May, July, September). We also calculate the percentage of generalist taxa present each month to determine if there is a ‘core’ macroinvertebrate community in the landscape, and we additionally hypothesize that streams exposed to more agricultural pressures will become more homogenous in community composition relative to less-impacted streams due to environmental filtering. We therefore expect that the local contribution to beta diversity (LCBD) will be lower in streams with a higher percentage of cropland in the catchment. Ultimately, we expect all three components (local factors, landscape factors, temporal factors) will likely interact to influence stream macroinvertebrate communities and that the incorporation of DNA metabarcoding will allow us greater resolution to detect complex patterns.

## 2. Methods

### 2.1 Stream monitoring and benthic macroinvertebrate collection

We have previously described our benthic macroinvertebrate collection and subsequent laboratory protocols for DNA metabarcoding in Gleason et al. (2022). Our study region encompassed six sub-watersheds in southern Ontario in the Lake Erie drainage basin. In total, we sampled twenty streams for aquatic macroinvertebrates, and recorded *in situ* water chemistry and physical habitat parameters, across three time points (May, July, and September 2019; Figure 1). Three sub-watersheds contained ‘singleton’ stream sites, while the remaining 17 streams were concentrated in the remaining three sub-watersheds (Figure 1). To be included in our field study, streams needed to be both wadeable and wet year long.

**Figure 1:**
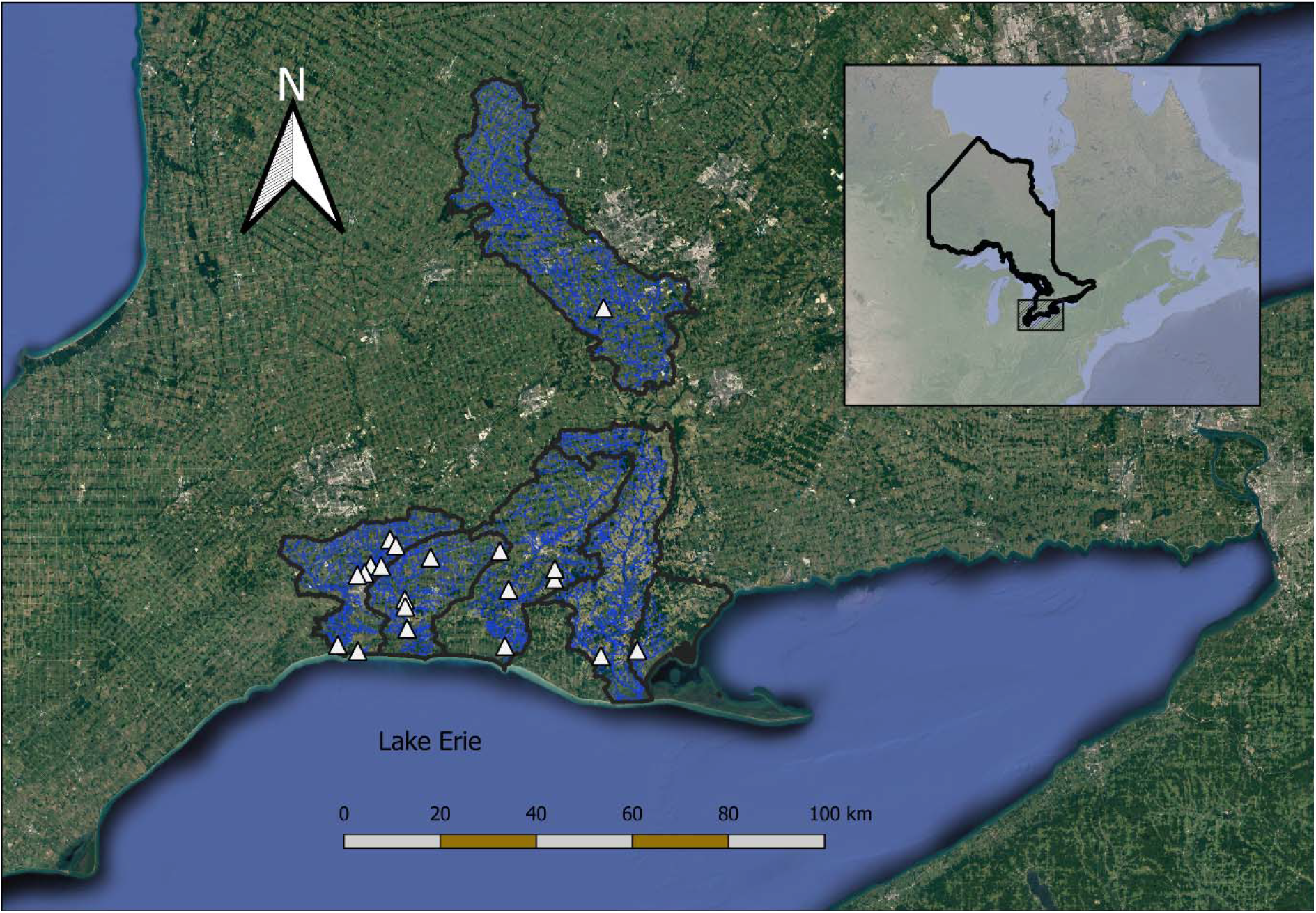
A map of the study region in southwestern Ontario, Canada. Sampling sites are represented by white triangles and their respective sub-watersheds are outlined in black.

Before disturbing the benthic sediment, we used an EXO2 Multiparameter Sonde (YSI Inc.) to measure *in situ* water chemistry (conductivity, total dissolved solids, dissolved oxygen, pH, dissolved organic matter). While depth, hydraulic head, and wetted width were also measured, they were not incorporated into data analysis due to the variability of such metrics in a single reach and because they are influenced by precipitation events (see Krynak & Yates, 2018). We measured riparian vegetation buffer width, stream valley slope, and stream sinuosity using Google Earth Pro (Google Inc.). For both buffer width and change in elevation between buffer and stream bed (bank slope), we averaged values between both sides of the stream and measured at five distances from the sampling site (500 meter intervals between 0 and two kilometers) before again averaging into a single representative value per site. We calculated stream sinuosity by dividing the five-kilometer within-stream distance by the straight-line distance between these points. To determine the percentage of land used for agriculture surrounding a stream, we first used the Ontario Flow Assessment Tool (OFAT; Ontario Ministry of Natural Resources and Forestry, 2020) to determine stream catchment boundaries in ArcGIS v. 10.6.1 (Esri, 2020) and then overlaid data from the Ontario Land Cover Compilation v. 2.0 (Ontario Ministry of Natural Resources and Forestry, 2016). We also used OFAT to determine stream order.

We collected four macroinvertebrate samples at each stream by choosing four transect locations ten meters apart across all microhabitats present (e.g., riffles, pools, differing sediments). Each sample consisted of a 3-minute kick sweep using a 500 μm D-net and was preserved separately in 95% ethanol on site. We cleaned all our sampling equipment (e.g., waders, nets, sieves, forceps) with 10% bleach and de-ionized (DI) water between each site. In total, there were 240 bulk invertebrate samples over our sampling period.

### 2.2 Laboratory work

We sorted benthic macroinvertebrates (arthropods, molluscs, and annelids) from sample debris under a dissection microscope and placed them in a sterile 20 mL tubes containing ten 4 mm diameter steel beads. We air dried samples and then homogenized them using an IKA Tube Mill (IKA, Staufen, Germany) at maximum speed (4000 rpm) for 15 minutes. We subsampled 20 mg (± 1 mg) of this homogenized invertebrate tissue powder into a sterile 2 mL tube and extracted DNA using a DNeasy Blood & Tissue Kit (manufacturer’s guidelines; Qiagen, Hilden, Germany). Following extraction, we quantified DNA concentration using a Qubit 3.0 Fluorometer (ThermoFisher Scientific, MA, USA) and stored the extracts in a −80°C freezer until PCR.

We used a two-step PCR protocol, first to amplify our marker, a 421 base pair fragment of mitochondrial cytochrome *c* oxidase subunit I (COI – the animal DNA barcode region; Hebert et al., 2003), and second to attach indexing primers for sequencing library preparation (see Gleason et al. 2022). We selected a degenerate primer pair (BF2 + BR2; Elbrecht & Leese, 2017), which is useful in community analysis to amplify a large range of invertebrate taxa, and has been successful with previous benthic samples collected from our study region (Gleason et al., 2021; Persaud et al., 2021). Our first PCR reaction consisted of 2.5 μL DNA extract, 12.5 μL of 2x master mix (Qiagen multiplex PCR kit; Qiagen, Hilden, Germany), 9 μL of molecular-grade water, and 0.5 μL of each primer (BF2 + BR2, initial concentration of 0.2 μM) for a reaction volume of 25 μL, and our thermocycling protocol was as follows: a 95°C initial denaturation for fifteen minutes; followed by 25 cycles of 94°C for 30 seconds, 50°C for 90 seconds, 72°C for 60 seconds; and a final extension at 72°C for ten minutes. We visualized the PCR product on precast 2.0% agarose e-gels (E-Gel 96 SYBR Safe DNA stain; ThermoFisher Scientific, MA, USA) to check for a band and purified PCR products using NucleoMag NGS clean up and size select magnetic beads (Macherey-Nagel, USA) at a 0.8x ratio (Milián-García et al. 2021). The second PCR was a 50 μL volume reaction including 5 μL of our purified PCR product, 25 μL of 2x Qiagen master mix, 10 μL of molecular water, and 5 μL each of the forward and reverse indexing primers (initial concentration 10 μM). The indexed library was again purified using NucleoMag beads (0.6x ratio), and the final product was visualized using e-gels. Each 96-well plate included eighty samples, eight negative controls (split between extraction, PCR, and sequencing), and eight technical replicates for a subset of samples to confirm consistency. We submitted the final libraries for sequencing on an Illumina MiSeq platform (V3 600 cycle kit) with 10% PhiX spike in at the Advanced Analysis Center at the University of Guelph on four separate runs.

### 2.3 Bioinformatics and quality control

We used the bioinformatics platform JAMP v. 0.67 (http://github.com/VascoElbrecht/JAMP) to process the raw sequences and have described similar pipelines and quality control in Gleason et al. 2022. The raw sequences were paired-end merged using USEARCH (v. 11.0.6668; Edgar, 2010), and cutadapt (v. 1.15; Martin, 2011) was used to trim the primer sequences from the ends of the reads. We filtered out reads that did not match our target sequence length (421 ± 10 bp) and removed low-quality sequences with an expected error (EE) score lower than 1. We selected a 97% clustering threshold and clustered OTUs using VSEARCH v. 2.18.0 (Rognes et al., 2016), and very low abundance OTUs (< 0.01% abundance across all samples) were removed. To match OTUs with taxonomic information, we used the Python program BOLDigger (Buchner & Leese, 2020) to match OTU sequences to the Barcode of Life Database (BOLD; Ratnasingham & Hebert, 2007).

All of our below data management and statistical analyses were performed using R v. 4.0.3 (R Core Team, 2021). To assess the quality of our data (e.g., checking for contamination and assessing sequencing depth), we used the R package metabaR v. 1.0.0 (Zinger et al., 2021). This package flags an OTU sequence as a contaminant if its relative abundance is highest in a negative control, and if more than 10% of reads of a sample corresponded to a contaminated sequence it would be considered a contaminated sample. No sample was flagged as containing contaminants, but there was a small number of reads in the sequencing controls, indicating tag-jumps had likely occurred during sequencing. To control for the influence of tag-jumps in the dataset, we selected a 0.001% abundance filter threshold (i.e., an OTU representing a relative abundance of less than 0.001% of total sequences of that OTU was converted to a zero in affected samples; see Gleason et al. 2022; Zinger et al., 2021). We only included benthic macroinvertebrate taxa in our dataset (arthropods, molluscs, annelids) and required a minimum match similarity threshold of 90% to the reference database (BOLD). Further details on our quality control pipeline and additional figures are available online as supplementary material in Gleason et al., 2022 (https://www.biorxiv.org/content/10.1101/2022.02.28.481642v1.supplementary-material). To create a single representative sample per stream, we averaged sequence reads across the four biological replicates. As sequence reads are not a reliable indicator of abundance due to biomass differences and primer biases (Elbrecht & Leese, 2015), we converted our final OTU table to a presence/absence matrix for downstream data analysis. Our final dataset consisted of three seasonal data frames (May, July, September) for twenty streams.

### 2.4 Statistical analysis

We calculated the monthly OTU richness at each site and used a repeated measures ANOVA to test for significant changes in overall richness between seasons (Figure 2). To determine the number of generalist taxa present in the dataset, we calculated how many OTUs were present in 14 or more (70%) of the 20 streams as well as what percentage of the dataset they made up. We also report the most frequently occurring OTU(s) each month and how many sites they were present at. Similarly, we calculated the proportion of rare OTUs that were present only in one site for each sampling month.

**Figure 2:**
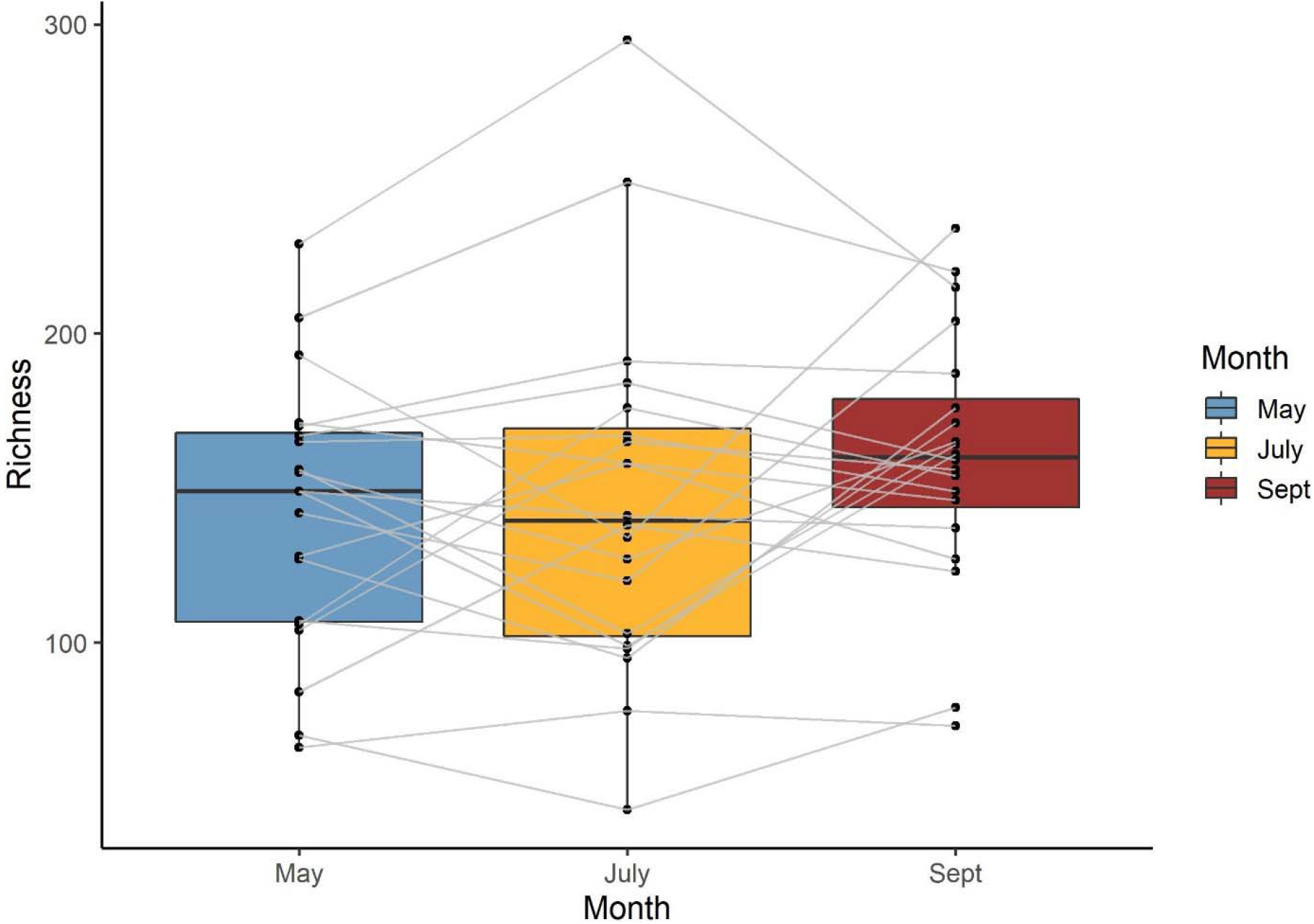
The OTU richness at each stream per month (n = 20). There is no significant difference between richness and sampling month (ANOVA: F_2,57_ = 0.76, *p* = 0.47). Grey lines represent the same site over time.

We used a perMANOVA to test if stream community composition remained consistent over time using the ‘adonis2’ function in the R package vegan (v. 2.5-7; Oksanen et al., 2020) with 999 permutations and a Jaccard dissimilarity matrix. We also used perMANOVA to test for the effects (and interaction) of the sampling month and the percentage of agricultural land use on the entire dataset. We then used the datasets subset by month to test the significance of agricultural land use on community composition for each monthly dataset, as well as the percentage of variation explained.

To reduce the number of water quality and local habitat variables in our ordinations while maximizing variation explained, we used forward selection prior to fitting RDA models (Blanchet et al.; Legendre & Gauthier, 2014). We used the ‘forward.sel’ function in the adespatial package (v. 0.3-14; Dray et al., 2021) with 999 permutations for both the entire dataset and each monthly dataset with the associated local metadata. We used a stopping criterion of alpha = 0.05 and an adjusted R^2^ equal to that of entire RDA model with all variables included for each monthly dataset respectively (0.168, 0.113, and 0.149). We selected significant environmental variables which met this threshold and only included these in downstream analyses for RDAs and variation partitioning. Similarly, we repeated this analysis for spatial parameters and network position (latitude, longitude, and stream order). For the entire dataset, the adjusted R^2^ for the entire model was zero, and thus no stopping criteria could be incorporated into the forward selection, and therefore only month was included as a variable in the RDA.

We performed a constrained ordination (dbRDA – distance-based redundancy analysis) using the ‘rda’ function in vegan on the entire dataset using sampling month, percentage of agricultural land use, and buffer width (see forward selection results) as constraints and a Jaccard dissimilarity matrix (Figure 3). We tested the significance of the model with an ANOVA with 999 permutations. We created the ordination plot using the R package ggplot2 v. 3.3.5 (Wickham et al., 2021) and plotted 95% confidence ellipses to delineate sampling months.

**Figure 3:**
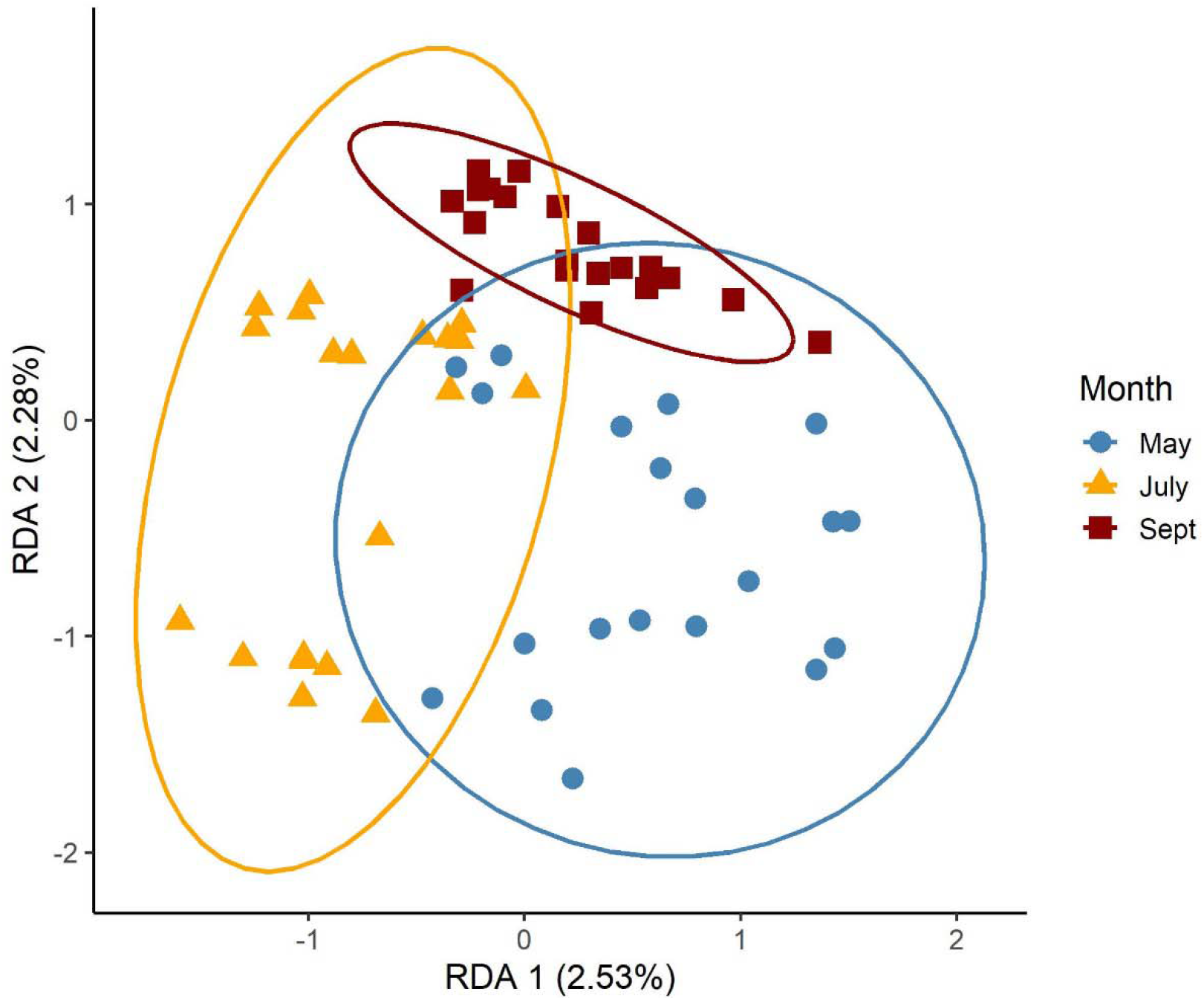
The plot of an RDA model with sampling month as a constraint (n = 60 streams). The percentage of variation explained by each axis is listed in brackets. Ellipses represent 95% confidence around sampling month. Symbols represent sites, and month is distinguished by colour and shape (May = blue circles, July = orange triangles, Sept = red squares).

We used the function ‘varpart’ from vegan to partition the variation explained by regional agriculture, local habitat parameters (those retained from forward selection), and spatial position (latitude, longitude, and stream order). The analysis was repeated for each month using a Jaccard dissimilarity matrix, and the proportions of variation explained (and their interactions) were visualized using Venn diagrams (Figure 4).

**Figure 4:**
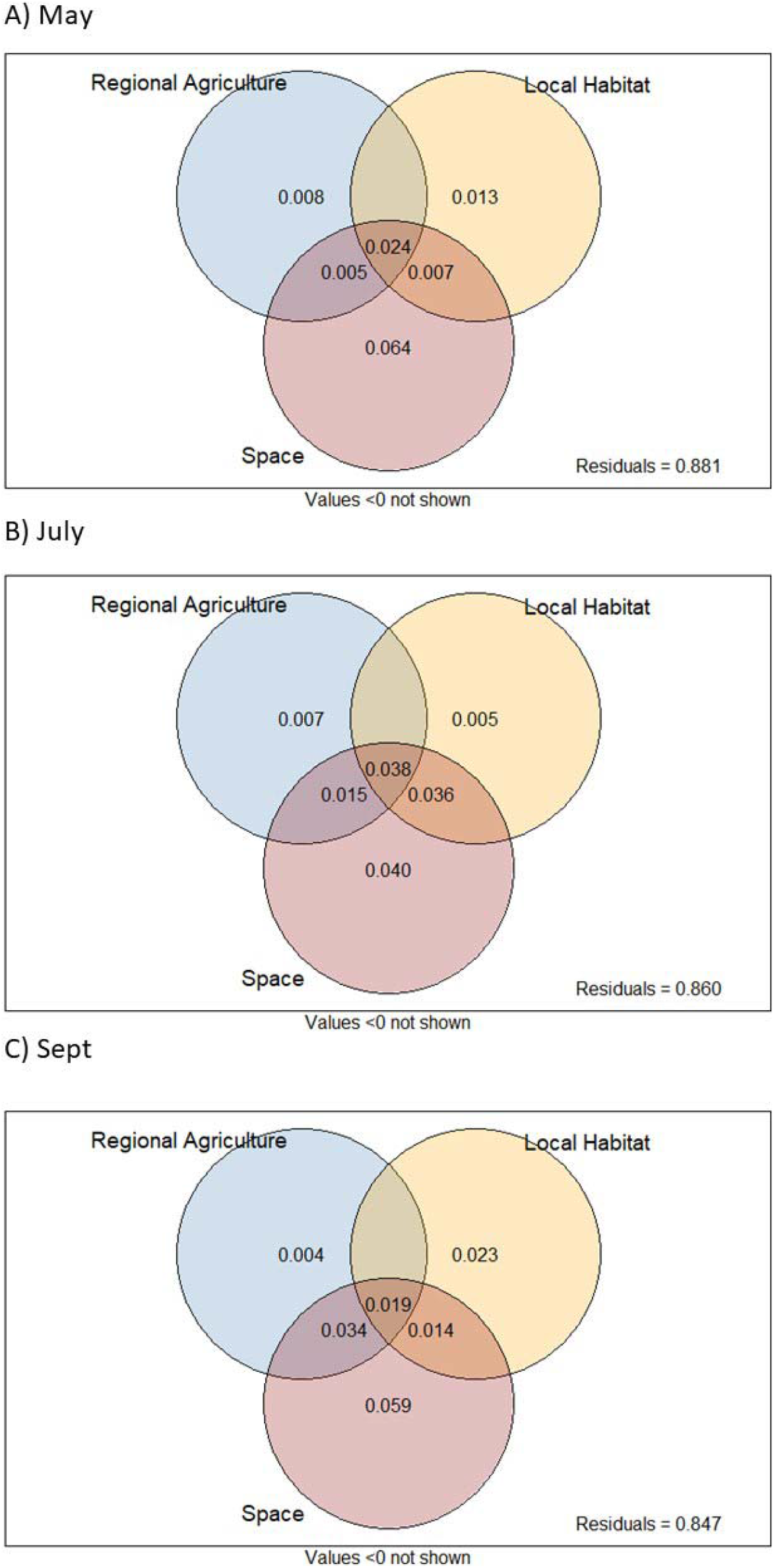
The results of variation partitioning visualized using Venn diagrams (circles not proportional to values) for each sampling month (May, July, September). The variables are Regional Agriculture (percentage of agricultural land use surrounding the stream), Local Habitat (the results of forward selection on a number of water quality and physical habitat parameters), and Space (latitude, longitude, and stream order). The residuals are shown in the bottom right corner of each plot.

We performed RDAs for each sampling month with the percentage of agricultural land use, the relevant local parameters selected from the forward selection analysis and spatial components (latitude, longitude, and stream order) as constraints. We tested the significance of each model separately using ANOVAs with 999 permutations. To visualize the effect of the selected habitat and spatial parameters on community composition, we plotted each monthly ordination using the first two RDA axes (Figure 5) and overlayed the relevant variables incorporated in the model to visualize any patterns in these parameters.

**Figure 5:**
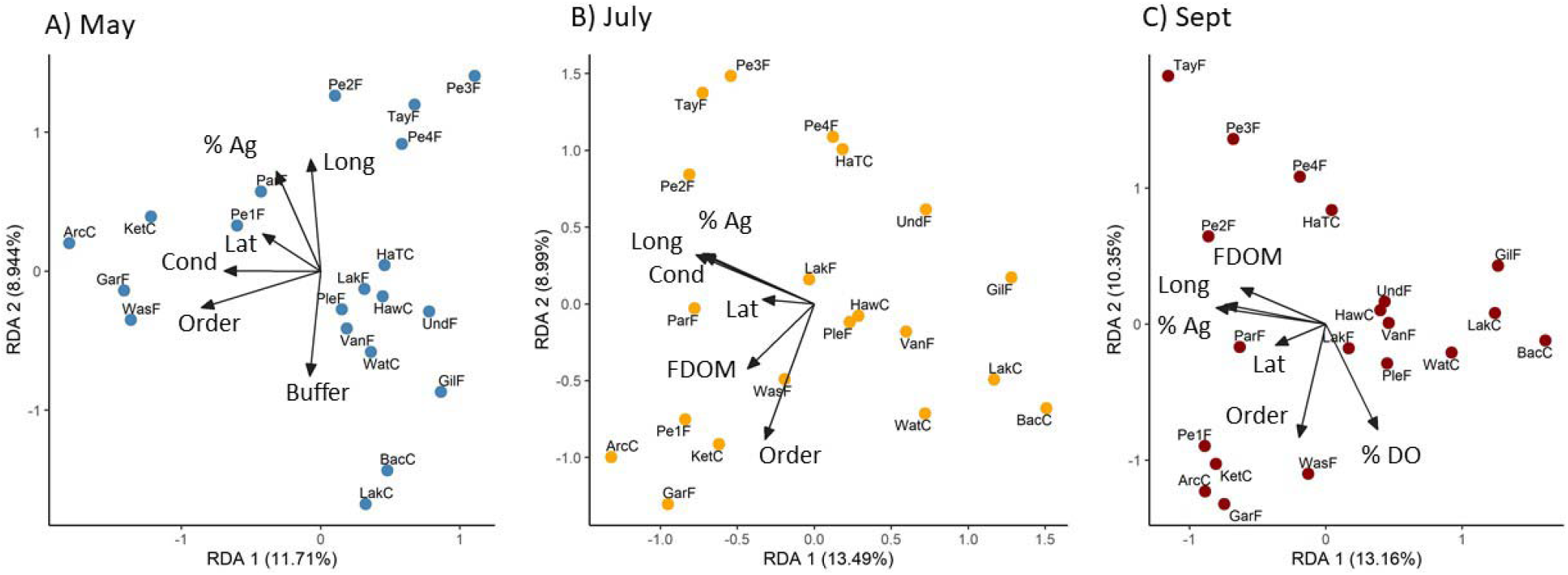
The RDA plots with sampling month, relevant local parameters, and the percentage of agricultural land use as constraints (n = 20 streams) by sampling month. The percentage of variation explained by each axis is listed in brackets. Symbols represent sites, and relevant habitat parameters are overlayed as vectors (% Ag = percentage of surrounding agricultural land use, Buffer = riparian buffer width, Cond = conductivity, % DO = percent dissolved oxygen, FDOM = fluorescent dissolved organic matter, Lat = latitude, Long = longitude, Order = stream order).

To determine whether streams became more homogeneous as agricultural land use increased, we calculated the local contribution to beta diversity (LCBD) using the ‘beta.div’ function in the adespatial package on a Jaccard dissimilarity matrix (Dray et al., 2021). LCBD is a measurement of how much a site contributes to the overall beta diversity of the group of sites, and the value is comparable between calculations (Legendre & Gauthier, 2014). A lower-than-average value indicates the site contributed less to the overall beta diversity (e.g., more homogeneous relative to other sites). To test for significant correlations between the percentage of agricultural land use and LCBD, we performed linear regressions for each sampling month.

## 3. Results

### 3.1 OTU richness and rarity

There were a total 41,978,040 sequences and 1681 OTUs in the final dataset post-bioinformatics and data cleaning. The majority of the OTU pool was made up of arthropods (1461), followed by 214 annelids and 21 molluscs. The total richness and average richness per site were consistent over the three sampling months, with 1076 OTUs in May (average of 148.6 ± 58.3 standard deviation), 967 OTUs in July (144.8 ± 45.7 SD), and 1045 OTUs in September (162 ± 42.2 SD), with no significant change in richness over time (Figure 2; repeated measures ANOVA: F_2,38_ = 2.08, *p* = 0.139).

For each month, the percentage of generalist taxa (present in 14 or more sites) was 1.02%, 0.62%, and 1.15%, respectively (Table 1). There were no OTUs that were present in every site during any month. In May, there was one OTU present in 19 out of the 20 sites (OTU 5 – *Orthocladius sp*.), in July the most frequent OTU occurred in 17 sites (OTU 29 – Naididae family), and in September there were two OTUs present in 17 sites (OTU 16 – *Hydropsyche betteni* and OTU 29 - Naididae). The proportion of rare OTUs (present in only one site) was consistent between sampling months at 46.7%, 44.4%, and 44.2%, respectively.

**Table 1:**
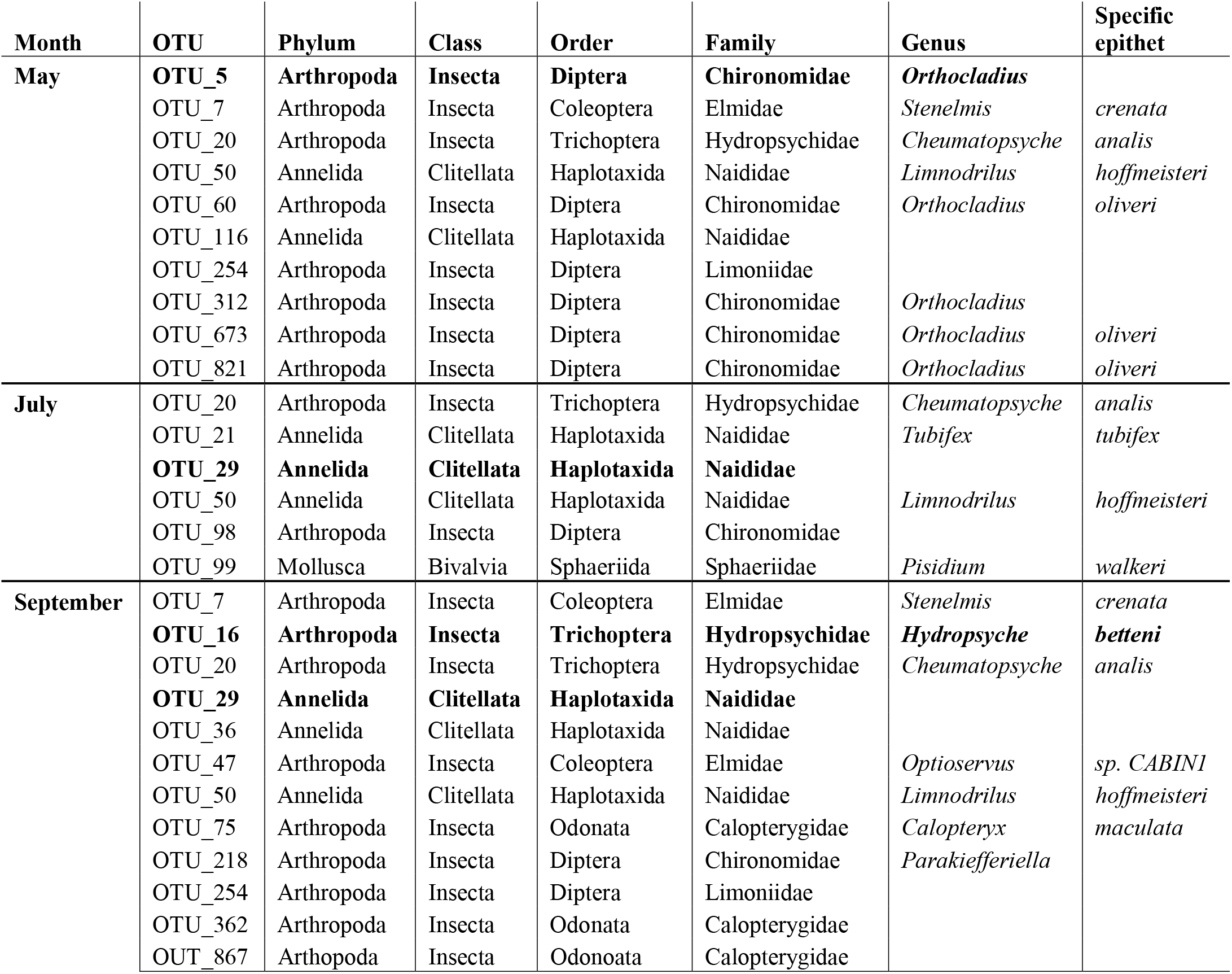
The number of generalist OTUs (present in at least 70% of streams) was calculated for each sampling month (May, July, September), and their proportion of the total monthly richness was 1.04%, 0.63%, and 1.15%, respectively. The OTU identification number is provided here as well as the taxonomic data that the OTU sequence matched to on the BOLD reference data (> 90% similarity). Bolded OTUs are those which occurred most frequently in that sampling month.

### 3.2 perMANOVA and forward selection of variables

The results of the first perMANOVA indicate that there is no significant variation explained by stream site over time (F_19,59_ = 0.87, R^2^ = 0.29, *p* = 0.99). In the second perMANOVA, there was a significant interaction of sampling month and the influence of agriculture (F_2,59_ = 1.27, R^2^ = 0.04, *p* = 0.04). The percentage of agriculture was not a significant factor independently for the entire dataset (F_1,59_ = 0.89, R^2^ = 0.01, *p* = 0.66), while month did explain significant variation in community composition (F_2,59_ = 1.53, R^2^ = 0.05, *p* = 0.005).

In the forward selection of environmental variables for the monthly datasets, conductivity (F = 1.58, *p* = 0.01) and buffer width (F = 1.48, *p* = 0.02) were selected for May (adjusted cumulative R^2^ = 0.05); conductivity (F = 2.08, *p* = 0.001) and dissolved organic matter (F = 1.43, *p* = 0.047) were selected for July (adjusted cumulative R^2^ = 0.07); and dissolved oxygen (F = 1.79, *p* = 0.007) and dissolved organic matter (F = 1.46, *p* = 0.05) were selected for September (adjusted cumulative R^2^ = 0.06). Spatial variables (latitude, longitude, stream order) were all significant for each monthly dataset (May adjusted cumulative R^2^ = 0.12; July adjusted cumulative R^2^ = 0.13; September adjusted cumulative R^2^ = 0.13).

### 3.3 Redundancy analysis and variation partitioning

The first RDA model using the entire dataset and month as an explanatory variable was significant (F_2,57_ = 1.44, *p* < 0.005), and the first two axes explained 2.53% and 2.28%, respectively (Figure 3). There were distinct clusters in stream by month, though September overlapped almost entirely with May and July.

The variation partitioning explained between 12-16% of community variation depending on the sampling month, with September having the highest proportion of variation explained (Figure 4). For every sampling month, the spatial component explained the most variation and overlapped with both agricultural land use and local habitat variables. All three variables had shared variation represented by the models in every month.

All monthly RDA models incorporating regional agricultural, local habitat quality, and spatial position were significant (May F_6,13_ = 1.53, *p* = 0.001; July F_6,13_ =1.51, *p* = 0.001; Sept F_6,13_ =1.60, *p* = 0.001; Figure 5). In May, the first axis represented 11.71% of the total variation and appeared driven by a correlation between higher stream order and conductivity (Figure 4–5A). The second axis of the May RDA (8.94%) aligned with spatial and agricultural parameters, with the percentage of agricultural activity and the riparian buffer width as opposing vectors (Figure 5A). In July, the first two axes represented 13.49% and 8.99% of the total variation, respectively (Figure 5B). The first axis appeared linked to stream order, while the second was a combination of spatial, agricultural, and local factors (Figure 5B). In September, the first axis explained 13.16% of the variation and aligned with stream order and the percentage of dissolved oxygen, while the second axis explained 10.35% and appeared to be a combination of spatial and agricultural parameters, along with dissolved organic matter (Figure 5C).

### 3.4 Local contribution to beta diversity

In May, there was a strong negative relationship between the percentage of agricultural land use and LCBD values, with sites surrounded by more agricultural land contributing less to overall beta diversity (Figure 6; adjusted R^2^ = 0.19, F_1,18_ = 5.44, *p* = 0.03). While a very weak negative relationship persisted in July, there was no longer any significant correlation between agriculture and LCBD (Figure 6; adjusted R^2^ = 0.02, F_1,18_ = 1.39, *p* = 0.25). There was a very weak positive relationship between agriculture and LCBD in September, but this was not significant (Figure 6; adjusted R^2^ = 0.01, F_1,18_ = 1.16, *p* = 0.30).

**Figure 6:**
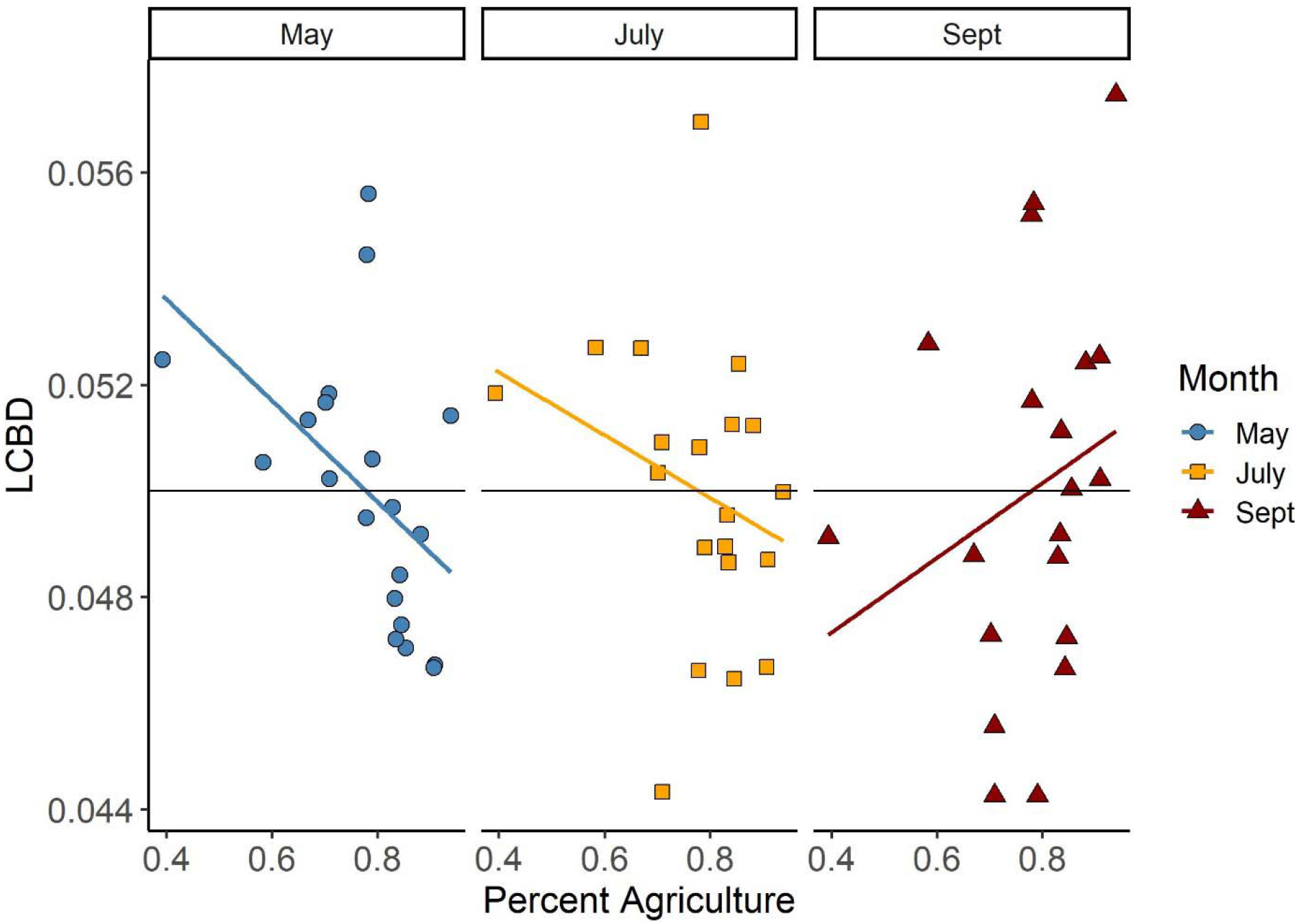
The percentage of agricultural land use in a stream catchment and the stream LCBD (local contribution to beta diversity) based on a Jaccard dissimilarity matrix for each sampling month (n = 20 streams). There was a significant negative relationship between agricultural land use and LCBD in May (adjusted R^2^ = 0.19, F_1,18_ = 5.44, *p* = 0.03) while July and September did not have significant correlations. The horizontal line represents the null value of 0.05, which would be the value if each site contributed equally to beta diversity (e.g., 1 divided by 20 sites).

## 4. Discussion

### 4.1 Environmental filtering and spatial components

A major goal of this work was to incorporate DNA metabarcoding into a metacommunity framework and demonstrate the utility of these two fields to monitor the effect of land use on stream ecosystems. Through variation partitioning and distance-based redundancy analyses, we demonstrate the relative influence of local and regional environmental conditions, as well as spatial position, on the community composition of aquatic macroinvertebrates. We observed that roughly twelve to sixteen percent of the variation in community composition was explained by a combination of 1) local environmental factors, such as water chemistry and riparian buffer width; 2) the percentage of agricultural land use in the drainage area; and 3) the spatial position of the stream. There was an overlap in the variation explained for almost every factor, suggesting that the spatial position of streams and both local and regional factors interact to influence community composition, and that it is challenging to disentangle these effects from one another. While the spatial components explained slightly more variation than the environmental variables included in our analyses, these values did not suggest that dispersal-based processes were more important than environmental filtering.

In a large meta-analysis of 95 datasets, Heino et al. (2015) observed that both spatial and environmental variables were typically poor predictors of how stream insect metacommunities were structured. Streams are very heterogenous habitats and support many rare species, which can make it challenging to describe patterns in invertebrate distributions (Heino et al., 2015). Previous research using DNA metabarcoding in Ontario streams supported this description of heterogenous streams as aquatic macroinvertebrate communities were highly variable within a single stretch and largely composed of rare OTUs (Gleason et al. 2022). However, while Heino et al. (2015) detected only weak relationships between environmental and spatial factors and macroinvertebrate community structure at genus-level resolution, we found significant relationships between all variables (local, regional, spatial) even though DNA metabarcoding increases the number of rare taxa detected.

While we found that immediate habitat conditions, regional land use, and spatial position all influenced the community composition of stream macroinvertebrates, the literature describing metacommunity dynamics in these systems is complicated, with varied results. Benthic macroinvertebrate communities in headwater streams in Maryland, USA, were primarily structured by local factors with no distance-decay relationship (i.e., species sorting with sufficient dispersal), whereas mainstem regions were influenced by a combination of both environmental filtering and dispersal effects (attributed to mass effects; Brown & Swan, 2010). While previous stream macroinvertebrate studies have concurred that the environmental component is the most important structuring factor in headwater steams (Göthe et al., 2013; Heino et al., 2012), we did not find that environmental filtering was the strongest component of stream community assembly in our study. Rather, we observed spatial components often explained more variation, though longitude was always along the same ordination axis as the percentage of agricultural land use, suggesting that there is a correlation between these variables. Our results suggest that land use does act as an environmental filter, though it is linked to the spatial location of the stream, which indicates an effect of dispersal limitation in these communities.

The majority of stream macroinvertebrate metacommunity studies are completed at mixed taxonomic resolution (often genus), and future work incorporating DNA metabarcoding is needed to draw stronger conclusions regarding niche and neutral processes. The spatial grain of a metacommunity ecology study is also an important consideration when interpreting results as the relative importance of environmental factors have been demonstrated to decrease at smaller spatial scales (Cottenie, 2005; Viana & Chase, 2019). Environmental processes have been described as more important for stream insects across large spatial scales (Heino et al., 2017), suggesting that stochasticity may be more evident at very small scales. We observed very high turnover between streams, with few generalists and many rare taxa occurring in only one stream, and at even smaller spatial scales, Gleason et al. (2022), observed a similar trend between stream transects. The high degree of community variability in these streams and the modest amount of variation explained by environmental and spatial variables in our study suggest that individual habitat patches may provide unique resources while also being subject to stochastic colonization (i.e., an interaction of niche and neutral assembly).

While the identity of early colonizers on a patch may be due to chance dispersal (i.e., stochastic effects), the presence of certain taxa can act as an additional biotic filter lens (e.g., co-existence theory; Aiken & Navarrete, 2014; Chesson, 2000; HilleRisLambers et al., 2012). Though we did not test for any effect of interaction between taxa, both abiotic aspects (e.g., water chemistry) and biotic aspects (e.g., microhabitats created by vegetation, facilitation, competition, predation) contribute to the local environment and thus community assembly (Thakur & Wright, 2017). Species traits are also an important component as environmental filtering can affect taxa differently based on their dispersal capabilities (Cañedo-Argüelles et al., 2015; Li et al., 2021), and therefore habitat fragmentation by land use may be more of an issue for weak-dispersing taxa.

### 4.2 Seasonal influence of land use

Habitat conditions, which influence community composition, can vary over time, and seasonality is an important consideration for metacommunity dynamics. Throughout the year, seasons can alter habitat conditions in ways that increase habitat availability (e.g., resource pulses) or through imposing new environmental filters, and thus it is surprising that the temporal component of metacommunity dynamics is often neglected (Holyoak et al., 2020). Aquatic macroinvertebrate community structure can change seasonally due to variations in habitat condition, such as in a floodplain in China where environmental filtering effects became stronger over a season as habitats became more fragmented (Li et al., 2022). Land use can also have a seasonal effect on streams, and the influence of contaminants on water quality has been shown to be elevated during wetter periods (Shi et al., 2017; Zhang et al., 2021).

In our study system, we observed that the relative importance of local environmental conditions varied each month while the percentage of regional cropland and spatial parameters remained consistent. Notably, the width of the riparian buffer (a static measurement over a season) explained the most variation in May and was directly opposite from percentage of agriculture land use along the second ordination axis, suggesting a gradient of disturbance. Riparian buffers can mitigate the influence of agriculture on stream macroinvertebrates (Marques et al., 2021), and perhaps this is at its most significant early in the cropping season. The importance of water chemistry parameters (conductivity, dissolved organic matter, dissolved oxygen) varied between months, suggesting that environmental conditions are not static within a site. Aquatic insect communities change seasonally due to differences in hatching and emergence times, and do not remain consistent over a season. As expected, we observed unique aquatic macroinvertebrate community compositions between our three sampling months. In addition to changes in agriculture pressures over time, it is also possible that each of these seasonal communities have their own sensitivities to agricultural pressures.

We expected that increased agricultural pressures would homogenize aquatic invertebrate communities through environmental filtering and predicted that streams with higher percentages of surrounding agricultural land use would contribute less to the beta diversity of the region (i.e., lower LCBD values). We found that there was only a significant negative relationship between LCBD and land use in May, meaning our hypothesis was only correct for one sampling month. Previous work on aquatic macroinvertebrates has discovered that land use causes both taxonomic and functional homogenization (e.g., low beta diversity) and that highly impacted streams are composed of a smaller subset of tolerant taxa (Martins et al., 2017; Siqueira et al., 2015). However, there is a clear knowledge gap regarding the effect of seasonality on land use and beta diversity, along with how differences in taxonomic resolution (i.e., molecular identification) can change these patterns. Here, we use DNA metabarcoding to demonstrate that aquatic invertebrate communities may be more vulnerable to homogenization in the spring (May in Ontario) and that increased riparian buffer widths can mitigate this effect.

### 4.3 Taxonomic resolution and OTUs

The use of DNA metabarcoding allowed us to incorporate higher levels of aquatic macroinvertebrate biodiversity into metacommunity analyses, which may provide a more accurate representation of how environmental and spatial components interact to influence community composition. In a study of Ontario streams, Martin et al. (2016) demonstrated that the response of aquatic macroinvertebrates to environmental conditions differed at three taxonomic resolutions (family, genus, and DNA barcoding). While environmental variables explained more community variation than spatial factors across all taxonomic resolutions, the proportion of variation explained was greatest at family-level (Martin et al., 2016). While this might suggest family-level identification is ideal, the decrease in variables (e.g., species to family) statistically allows for more variation to be explained by decreasing the ‘noise’ of within-family patterns. It is also possible that some species within a genus, or even genera within a family, are interchangeable through neutral processes rather than subject to specific environmental tolerances. The theory of lumpy species distribution suggests a reconciliation between the niche and neutral approaches where very similar species do not directly compete (e.g., they become competitively neutral) but instead compete with other ‘clumps’ of likewise similar species (Scheffer & van Nes, 2006). Martin et al. (2016) observed that environmental covariates explained less community composition as taxonomic resolution decreased in coarseness, suggesting that species can be interchangeable within a genus under certain conditions.

A limitation of using molecular identification is that the use of OTUs as taxonomic units is not a guarantee of species-level identification, and it is possible that some species may be mistakenly lumped together or split across multiple OTUs. The choice of clustering threshold can influence the number of OTUs generated, as well as how closely these clusters match actual species delineations (Clare et al., 2016). We observed very high counts of OTUs from the dipteran family Chironomidae (nearly one third of our dataset), and the majority of these OTUs could not be assigned a binomial species name due to gaps in the reference database. Our results are consistent with other DNA metabarcoding studies, which similarly detected high levels of chironomid OTUs (e.g., Beermann et al., 2018), and future work establishing reference databases and species delineations are necessary for this large yet understudied group. For example, Lin et al. (2017) revealed that some chironomid groups had very high within-species divergences, and the lack of a ‘barcode gap’ can severely interfere with the effectiveness of OTU clustering (Čandek & Kuntner, 2015). An alternative to OTU clustering, which can overcome this issue, is the use of haplotype-level diversity (exact sequence variants or ESVs) to avoid clustering altogether (Porter & Hajibabaei, 2020), which may be an effective tool for highly diverse groups such as chironomids.

### 4.4 Conclusions and future work

The use of DNA metabarcoding to uncover large amounts of diversity in a system can provide greater insight into the relative importance of environmental filtering and stochasticity in freshwater ecosystems, particularly those impacted by anthropogenic land use. There is a clear need to move beyond using only local assessments to evaluate habitat conditions, as stream networks are large, connected ecosystems, which vary temporally (Cid et al., 2022; da Silva et al., 2021; Gounand et al., 2018). In this study, we demonstrate the importance of both environmental and spatial components in structuring aquatic invertebrate metacommunities and that these processes were also seasonally influenced. The temporal component of environmental variability and metacommunities is an area which demands future work, particularly as shifts in environmental timings are changing due to climate change (Holyoak et al., 2020). In addition to seasonal effects, we also observed very high diversity in our stream sites, with few generalist taxa and many rare species. In general, more research needs to be done globally to detect trends in freshwater insect biodiversity (Jähnig et al., 2021), ideally incorporating both taxonomic experts as well as molecular approaches. In this work, we have demonstrated the utility of DNA metabarcoding for improved resolution in biomonitoring programs and to detect patterns in metacommunity dynamics. Advances in these research directions should stress the utility of DNA metabarcoding and applied metacommunity ecology in conservation, and ideally improve our understanding of how best to protect freshwater resources.

## Data availability

Sequencing data is available publicly on NCBI’s SRA under accession PRJNA783201.

## Author contributions

JEG, RH, and KC conceived the ideas; JEG designed the study, performed the field and laboratory work, performed the bioinformatics and data analysis, and wrote the first draft of the manuscript; JEG, RH, and KC edited the manuscript.

## Acknowledgements

We would like to acknowledge that our stream sampling sites are located on the traditional territory of the Mississaugas of the Credit First Nations, the Anishnabek, Haudenosaunee (Iroquis), and Ojibway/Chippewa peoples.

We are grateful to Ian Thompson, Marie Gutgesell, Emily Champagne, and Christine Dulal-Whiteway for their assistance with field work. Dr. Yoamel Milián-García provided helpful advice regarding the molecular lab work pipelines, and Dr. Sarah (Sally) Adamowicz provided suggestions on an early draft of this manuscript. We would also like to thank staff at the following conservation authorities for facilitating sampling: Catfish Creek CA, Kettle Creek CA, Long Point CA, Nature Conservancy Canada and Thames Talbot Land Trust.

